# Infection of the *Biomphalaria glabrata vector snail* by *Schistosoma mansoni* parasites drives snail microbiota dysbiosis

**DOI:** 10.1101/386623

**Authors:** Anaïs Portet, Eve Toulza, Ana Lokmer, Camille Huot, David Duval, Richard Galinier, Benjamin Gourbal

## Abstract

Host-associated microbiota cari affect the fitness of its host i η a number of ways, including the modification of host-parasite interactions and thus the outcome of disease. *Biomphalaria glabrata* is the vector snail of the trematode *Schistosoma mansoni,* the agent of human schistosomiasis, causing hundreds of thousands of deaths every year. Here, we present the first study of the snail bacterial microbiota in response to *Schistosoma* infection. To examine the interplay between *B. glabrata, S. mansoni* and snail microbiota, snails were infected and the microbiota composition was analysed by massive 16S rDNA amplicon sequencing approach. We characterized the *Biomphalaria* bacterial microbiota at the individual level in both naive and infected snails. Sympatric and allopatric strains of parasites were used for infections and re­infections to analyse the modification or dysbiosis of snail microbiota in different host-parasite co-evolutionary contexts. Concomitantly, using RNAseq data, we investigated the link between bacterial microbiota dysbiosis and snail anti-microbial peptide immune response. This work paves the way for a better understanding of snail/schistosome interaction, and would have critical consequences in terms of snail control strategies for fighting schistosomiasis disease in the field.

## Introduction

Several studies have shown that the microbiota interacts with pathogens and/or host immunity in invertebrates. For example, the midgut microbiota of the mosquito *Aedes sp.* elicits a basal immune activity of the immune system (Ramirez et al., 2012) mostly by activating Toll pathways (Xi et al., 2008). In *Anopheles gambiae,* some members of its microbiota can limit malaria transmission, by inducing a wide antimicrobial immune response (Gendrin, 2017). More precisely, bacteria of the genus *Enterobacter,* found in the mosquito microbiota, were shown to produce reactive oxygen species (ROS) that inhibit *Plasmodium* development (Cirimotich et al., 2011). Furthermore, the endosymbiotic bacteria *Wolbachia* induce expression of immune genes, like TEP1 (Thioester-containing protein 1), LRIM1 (Leucine-Rich Immune Molecule 1) or defensin 1 (Joshi et al., 2017). In *Drosophila,* the bacterial microbiota is necessary to produce the Pvf2, a PDGF/VEGF-like growth factor that restricts enteric viral infection (Sansone et al., 2015). The microbiota may also play a role in immune priming. For example, in *Anopheles gambiae,* hemocyte priming during the *Plasmodium* infection is naturally induced by the gut microbiota, whose absence results in more severe infections and re-infections (Rodrigues et al., 2010). Similarly, the gut microbiota is necessary for immune priming in *Tribolium castaneum* and efficient response against *Bacillus thuringiensis for Tenebrionis* (Futo et al., 2015). Finally, host immunity can influence the tolerance or control the microbiota. In the tick *Ixodes scapularis,* the protein PIXR, secreted by the tick’s gut, inhibits the bacterial biofilm formation and supports gut eubiosis (healthy gut microbiota), its inactivation leading to dysbiosis. This change in gut microbiota facilitates the colonization by *Borrelia burgdorferi,* the Lyme disease agent (Narasimhan et al., 2017). In *Drosophila,* for example, the intestinal homeobox gene Caudal regulates the commensal-gut mutualism by repressing nuclear factor kappa B-dependent antimicrobial peptide genes (Ryu et al., 2008). In Hydra, the innate immune sensors and effectors protect not only against pathogens but also control microbiota homeostasis (Augustin et al., 2010). Moreover, host specific antimicrobial peptides (Fraune et al., 2010; Franzenburg et al., 2012) and a rich repertoire of pattern recognition receptors (Bosch, 2013) are involved in maintaining homeostasis between the host and the resident microbiota. Similarly, bactericidal permeability-increasing proteins (BPIs) shape bacterial communities in the light organ of the squid *Euprymna scolopes* and prevent their spill over to other tissues (Chen et al., 2017).

*Biomphalaria glabrata* is a freshwater snail (Lophotrochozoa, Planorbidae), living in inter­tropical regions of Latin America, in rivers, ponds, waterways and other freshwater environments. *B. glabrata* snails have important medical and epidemiological impacts due to their role as the main vector of *Schistosoma mansoni* (Lophotrochozoa, Platyhelminthes, Trematoda), the agent of intestinal schistosomiasis. Schistosomiasis is the second most widespread human parasitic disease after malaria, affecting over 200 million people worldwide and causing 200 000 deaths annually (WHO, 2002). The *Schistosoma* adult parasites mate in the human host venous system. Female worms produce eggs that cross endothelial mesenteric vessels and intestinal epithelium to reach faeces and finally the aquatic environment. Once in the water, eggs hatch and release miracidia, the free-living snail-infective parasite stage. At the following step of the life cycle, the miracidium needs to infect the freshwater snail *B. glabrata.* Intensive asexual multiplication in the snail tissues leads to the continuous production of hundreds of generations of cercariae, the free-living human-infective parasite stage.

At present, there is no effective vaccine against schistosomiasis and the treatment relies on a single chemotherapeutic treatment, the Praziquantel (Doenhoff et al., 2009), against which resistance has been observed (Fallon and Doenhoff, 1994). Molluscicides have been also used to impair *Schistosoma* transmission in the field. However, the dramatic effects of molluscicides on natural environments prompt us to seek new ways to prevent and / or control this disease in the field (Tennessen et al., 2015). We propose that a better understanding of ecological interactions between, *Schistosoma mansoni, Biomphalaria glabrata* snails and its associated microbiota might represent an interesting approach in search for alternative control strategies.

In this context, we have conducted numerous studies on immunological interactions between the snail and the parasite (Mitta et al., 2005; Bouchut et al., 2008; Moné et al., 2011; Coustau et al., 2015; Galinier et al., 2017; Mitta et al., 2017). We have demonstrated that the nature of the snail immune response depends on the strain/species of host or of parasite considered. Depending on both the host and the parasite intrinsic capacities, parasites either develop normally in snail tissue (compatible interaction) or are encapsulated by hemocytes (the snail immune cells) (incompatible interaction) (Mone et al., 2010; Mitta et al., 2012; Coustau et al., 2015) (Fig. 1). Depending on the ecological context, totally different immunobiological interactions have been described in *B. glabrata/S. mansoni* interactions. If immunosuppression occurs during a sympatric interaction, an activation of the immune response is observed during an allopatric interaction (Portet et al., 2019). Moreover, following primo-infection and successive challenges, a shift from a cellular immune response toward a humoral immune response has been described (Pinaud et al., 2016) (Fig. 1).

**Fig. 1:**
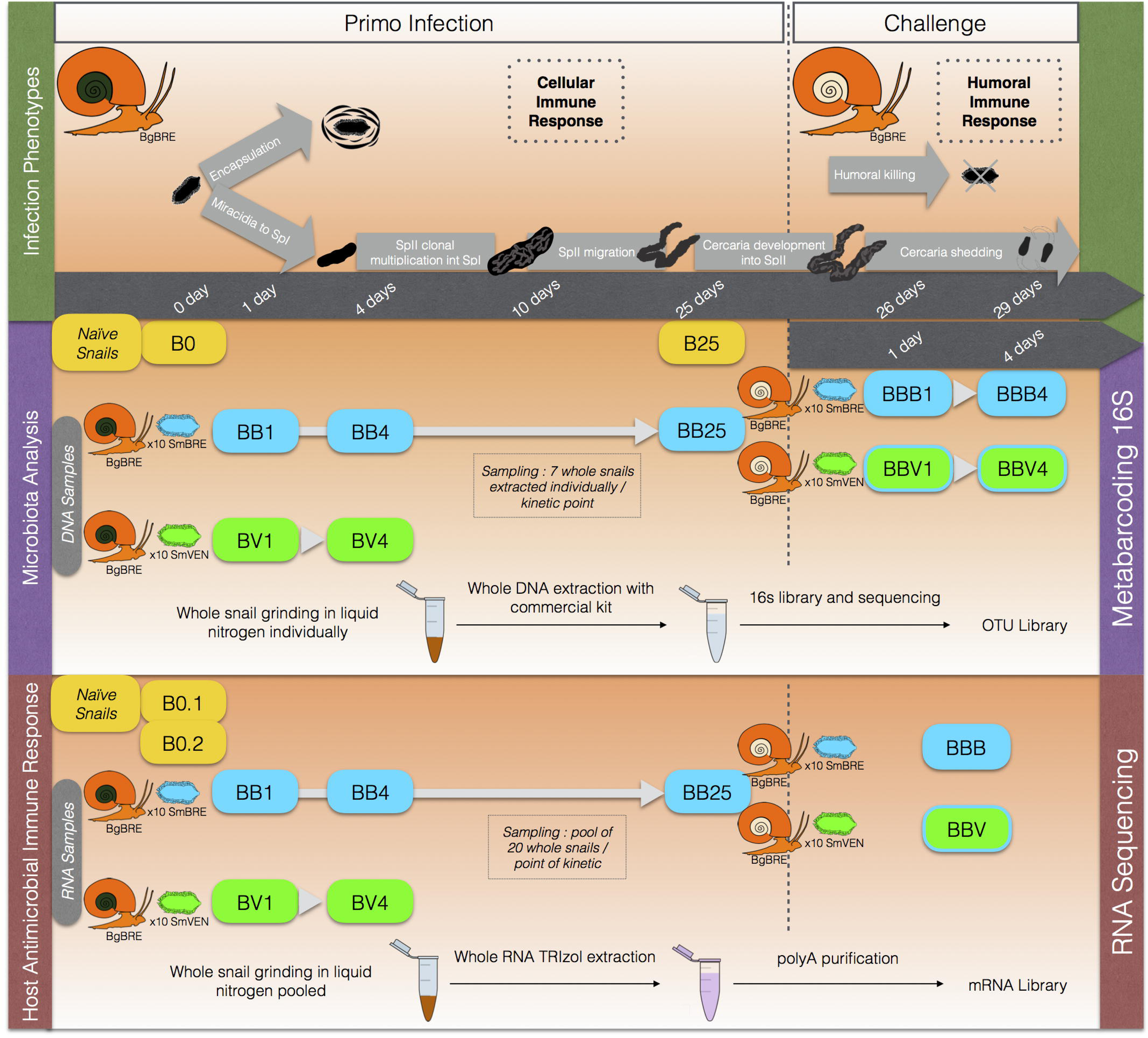
Experimental protocol.

However, in the present host-parasite model, only few studies examined the bacterial microbiota of *Biomphalaria glabrata* so far (Ducklow et al., 1979; Silva et al., 2013). The study of aerobic heterotrophic cultivable flora of 200 snails (Ducklow et al., 1979) revealed five predominant bacterial genera including *Pseudomonas, Acinetobacter, Aeromonas, Vibrio* and *Enterobacter.* I η a more recent study, the *Biomphalaria glabrata* bacterial microbiota has been characterized using 16S rRNA sequences (Silva et al., 2013) of cultured isolates, identifying six additional bacteria genera: *Citrobacter, Cupriavidus, Rhizobium, Stenotrophomonas, Klebsiella* and *Sphingomonas.* In addition, diversity and composition of bacterial microbiota differed between resistant and susceptible *Biomphalaria* phenotypes, with differences in relative abundances of *Gemmatimonas aurantiaca* and *Micavibrio aeruginosavorus.* This observation supports the potential link between the resistance against *Schistosoma* parasite and the structure of snail-associated microbiota (Allan et al., 2018). However, no studies have examined changes in bacterial microbiota of *Biomphalaria* snails by culture-independent methods nor following Schistosoma infections so far.

Taking into account all the peculiarities of this system, the aim of the present study was to investigate variation of microbiota in snail populations as well as to assess the influence of varying immune response against *Schistosoma* on the host microbiota composition and dynamics. To reach this goal, we first characterized the bacterial microbiota of naive *Biomphalaria glabrata* snails using 16S rDNA metabarcoding. Then, we analysed changes in microbiota composition following, (i) primo-infections with sympatric and allopatricS. *mansoni* parasite isolates displaying the same prevalence and intensity phenotypes and, (ii) subsequent secondary challenges with homologous or heterologous parasite strains. As snail immune response strongly differs between such diverse snail/parasite combinations, we concomitantly analysed the snail immune response using a massive sequencing transcriptomic approach (RNAseq) in order to link it with the observed changes in the composition and diversity of the microbiota.

## Results

### Characterization of healthy *B. glabrata* microbiota

To examine the stability of naive BgBRE snail microbiota, we first inspected the microbiota diversity and composition in the control naive snails at day 0 and 25 of the experiment. We found no significant differences between the B0 and B25 snails in any of the alpha diversity indices (Table S3, Table S4).

Naive snails showed little inter-individual variability and a stable composition at the phylum level over time (Fig. 2A, Table S3, Table S4), with Proteobacteria, Bacteroidetes, Cyanobacteria and Planctomycetes phyla being the most represented (Fig 2A). Moreover, the bacterial microbiota of naïve BgBRE snails displayed a considerable temporal and inter­individual stability at the family level (Fig. 2B). The stability of naïve snail microbiota composition over time (Fig. 2) could be related to the laboratory rearing conditions and laboratory environmental abiotic variables (water composition, temperature, pH, food) that were tidily controlled and that did thus not result in changes in host snail physiology or metabolism.

**Fig. 2:**
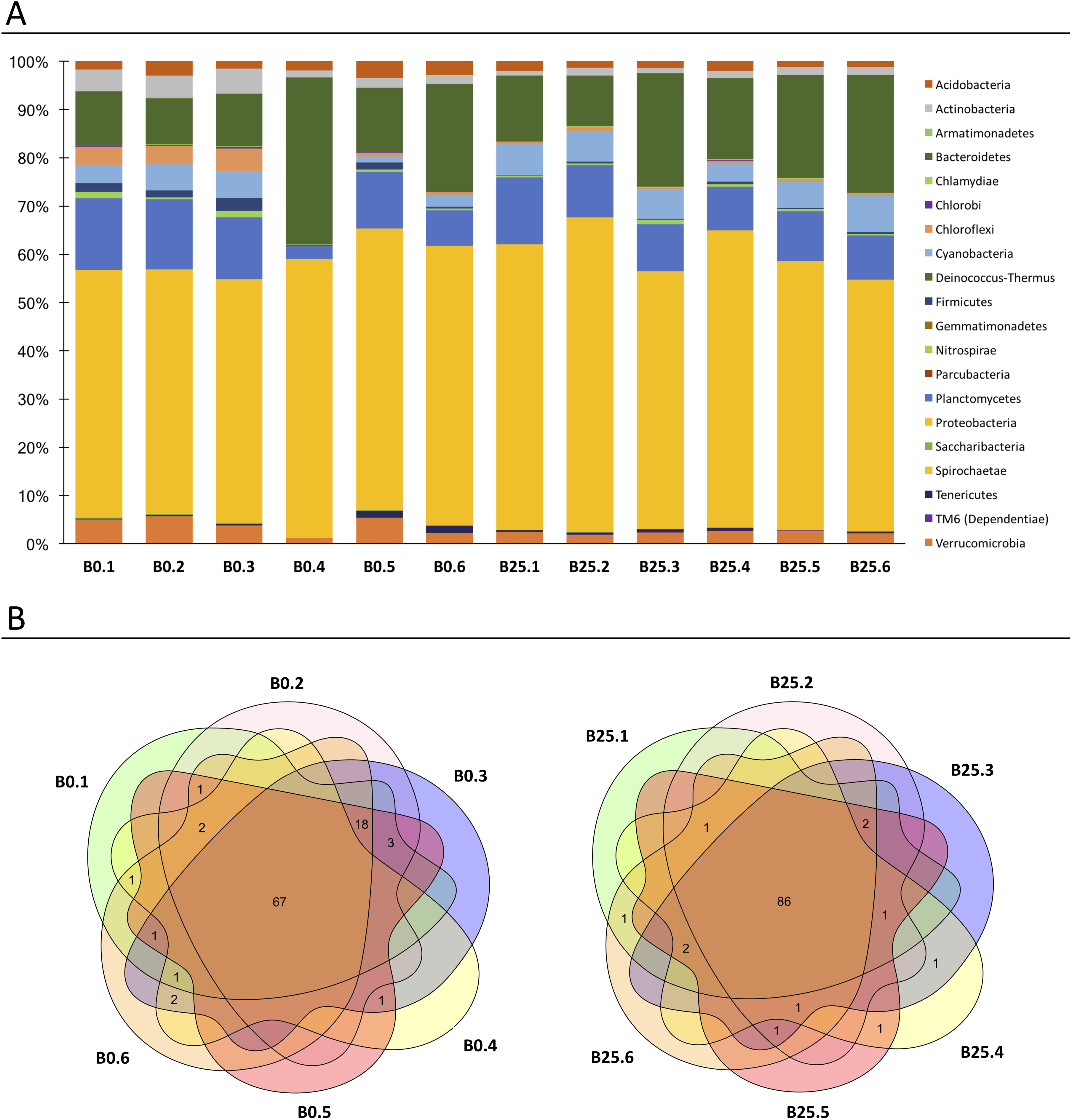
*Biomphalaria glabrata* microbiota characterization. Characterisation of the *Biomphalaria* bacterial microbiota of six naive snails recovered at the start of experimentation (B0.1; BO.2; BO.3; B0.4; BO.5 and BO.6) and 6 naive snails recovered 25 days after the start of experimentation (B25.1; B25.2; B25.3; B25.4; B25.5 and B25.6). A. Phylum level composition of the 20 most abundant OTUs phyla among the 12 naive snails. B. The Venn diagram represents the number of the 97 OTUs families which shared between the 6 naive snails at BO (left Venn diagram), and between the 6 naive snails at B25 (right Venn diagram).

In terms of composition, we observed that 67 (69%) and 86 (89%) out of 97 identified families were shared by all individuals of B0 or B25 naive snails respectively (Fig. 2B). Those results were used to determine the core microbiota. We defined core microbiota as the families that were present in 100% of the naive snails. Applying this definition, we identified 62 out of 97 families found in all naive individual snails (B0 and B25 altogether), and thus constituting the *B. glabrata* core-microbiota (Table S7 and S8).

### Microbiota dynamics following *B. glabrata* infections by *S. mansoni*

After establishing the stability of naive *Biomphalaria* bacterial microbiota, we investigated whether *Schistosoma mansoni* infections affected the microbiota structure and dynamics. To investigate the influence of parasite infections on the bacterial microbiota, we analysed microbiota dynamics following sympatric or allopatric primo-infections and homologous or heterologous challenges (Fig. 1).

We did not observe any significant changes in alpha diversity during the course of primo-infection compared to naive snails, excepted a decrease in the Shannon’s Η diversity index at the fourth day after sympatric infection (Fig. 3, Table S3, Mann-Whitney U test: Table S4). Conversely, all indices changed significantly following homologous or heterologous challenges (Fig. 3, Table S3, Table S4). Indeed, the observed species richness (Mann-Whitney U test, p=0.002), the Chao 1 richness index (Mann-Whitney U test, p=0.006), Shannon diversity index (p=0.0004) and Pielou evenness index (Mann-Whitney U test, p=0.009) were significantly reduced following homologous or heterologous challenges compared to naive and primo-infected snails (Fig. 3, Table S3, Table S4). However, the observed drop in alpha diversity disappeared by the fourth day after heterologous challenge (Fig. 3, Table S4). In addition, alpha diversity was mainly affected by the challenge, regardless of its type (homologous or heterologous). The primo-infection did not significantly affect alpha diversity except for the sympatric combination at day 4 after infection (Table S4).

**Fig. 3:**
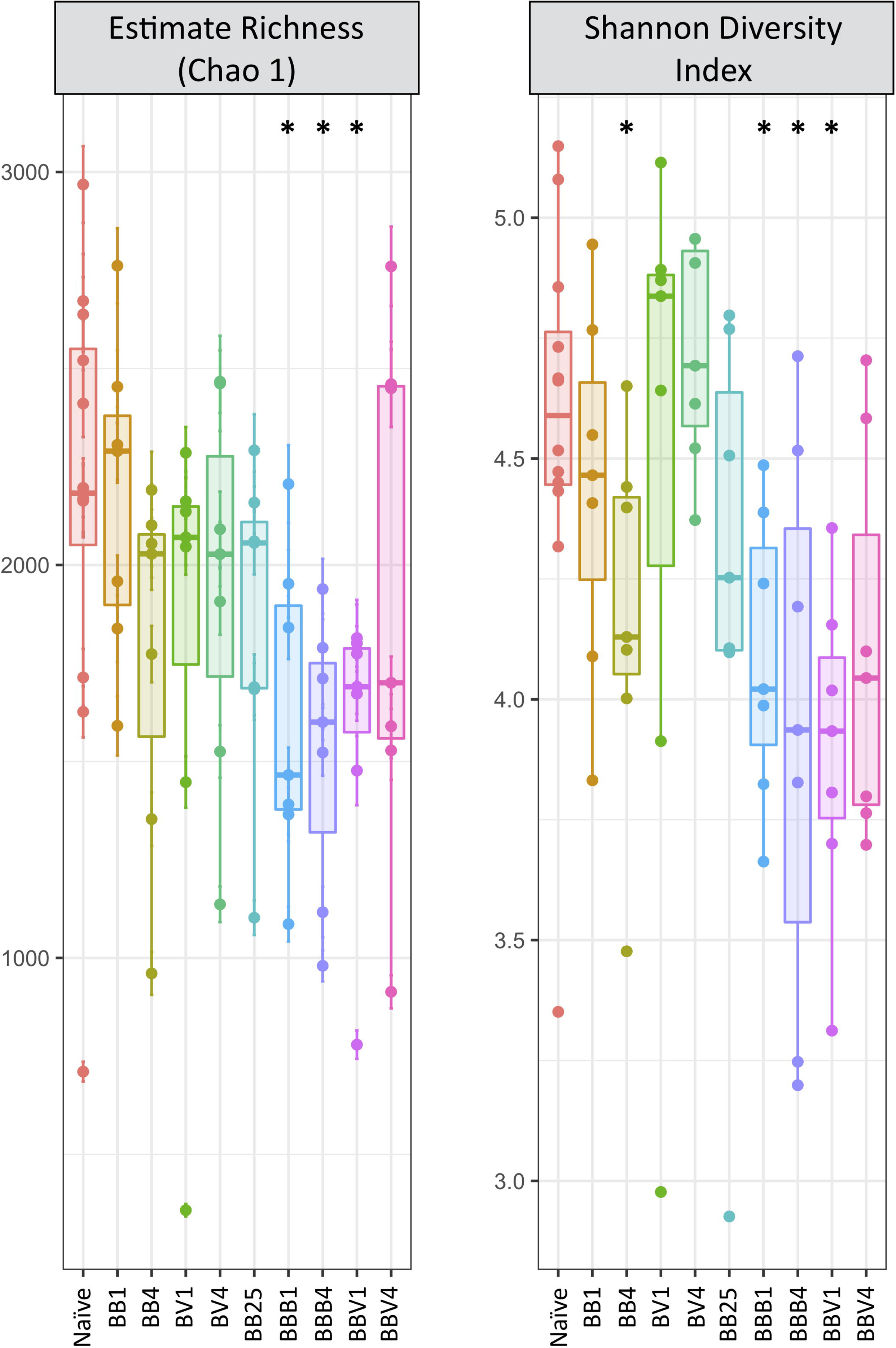
Microbiota alpha Diversity. Boxplots of Chaol and Shannon indices for all samples. For the Naive condition, B0 and B25 snails were pooled; BB: primo infection of BgBRE by SmBRE; BV: primo infection of BgBRE by SmVEN; BBB: primo infection of BgBRE by SmBRE and challenge by SmBRE; BBV: primo infection of BgBRE by SmBRE and challenge by SmVEN. The time point is mentioned with 1, 4 or 25 corresponding to the day after primo-infection or challenge. The differences between naive and infected conditions were tested with a Mann-Whitney U test and significant differences mentioned with “*”.

Regarding the beta diversity, Principal Coordinate Analysis (PCoA) of Bray-Curtis dissimilarities revealed that BB25 samples grouped together with the naive snail samples (B0 and B25) and were separated from the infected-snail samples along both axes (Fig. 4A, Table S5). However, the BB25 samples were significantly different from the naive snail samples (Table S5). The fact that, BB25 grouped with naive snails rather than with infected-snail samples suggests that the snail microbiota is resilient to infection, with a tendency to recover between day 4 and day 25 after the primo-infection (Fig. 4A). Analysis of Bray-Curtis dissimilarity index revealed a significant difference between naive and primo-infected samples (p=0.001) and also between naive and challenged samples (p=0.001) (Table S5). Concerning the infected-snail samples, all experimental samples were significantly different from each other with the exception of BB1 versus BB4 and BBB1 versus BBV1 (Table S5).

**Fig. 4:**
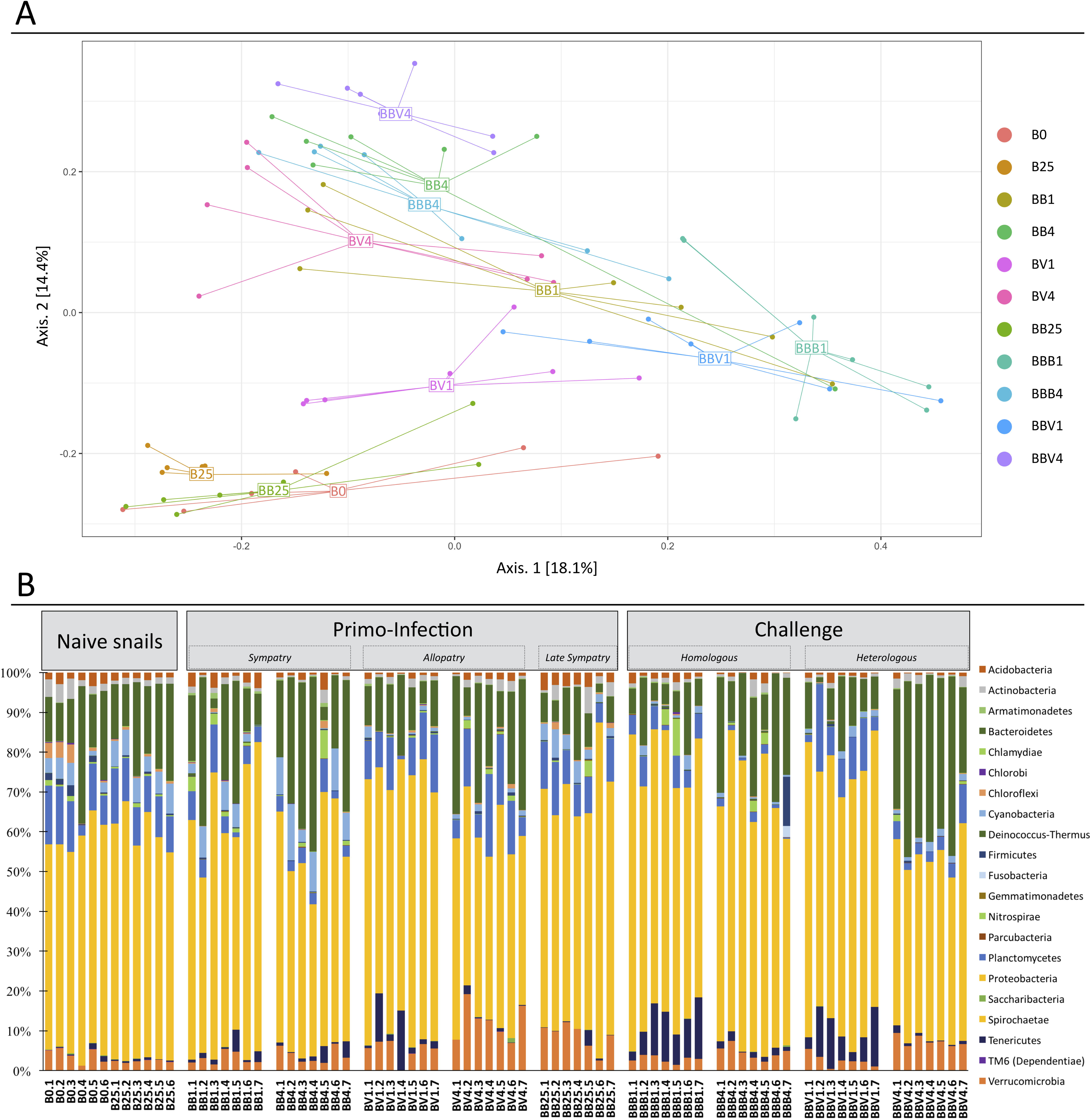
Beta diversity and bacterial communities following *Biomphalaria* infection. Dynamics of the bacterial microbiota of *Biomphalaria glabrata* following *Schistosoma* primo-infection and challenge. A. Functional diversity comparisons of *Biomphalaria* microbiota along infection. Principal coordinate analysis of pairwise Bray-Curtis distances between all infection type and time samples. Axes represent the two synthetic variables explaining the greatest proportion of variation in the data set. The sample name indicated in the figure corresponds to the centroid of all the biological replicates points of the respective experimental sample. B. Phylum level composition of the 20 most abundant OTUs among all points of the kinetic. In this representation, the replicate naive snails were pooled for more readability.

In addition, the second PcoA axis separated the samples according to the course of infection (i.e. the day 1 from the day 4 samples) (Fig. 4A), whereas the day 1 after-challenge (BBB1, BBV1), the day 1 after-primo-infection (BB1, BV1) and all of the day 4 infection samples (BB4, BV4, BBB4, BBV4) were separated along the first axis (Fig. 4A). Interestingly, day 1 after-challenge samples (BBB1, BBV1) were more different from the naive microbial communities than the day 1 primo-infection samples (BB1, BV1). Moreover, the second axis separated the sympatric primo-infection (BB1) from the allopatric primo-infection (BV1) (Fig. 4A). Finally, all day 4 samples grouped together reflecting the similarity between these samples regardless of experimental infection conditions (Fig. 4A). Even if the day of infection appears as the main explaining factor, these results indicate that the snail microbiota response to the infection also depends on the type of infection (sympatric vs. allopatric or primo-infection vs. challenge) and suggest the existence of predictable dynamics.

We further investigated the microbial community dynamics following infection, and we observed some consistent changes in response to experimental infection type (Fig. 4B). The relative abundance of Tenericutes increased after infection, particularly at day 1 after challenge (BB25 vs BBB1, student Τ test Ρ = 0.0114; BB25 vs BBV1 student Τ test Ρ = 0.0209). Increase in Verrucomicrobia following allopatric primo-infection (B0/B25 vs BV4, student Τ test Ρ = 6.53 e-06) and homologous / heterologous challenge (B0/B25 vs BBB4/BBV4, student Τ test Ρ = 0.0012) was most prominent at day 4 after infection (Fig. 4B). Interestingly, high relative abundance of Verrucomicrobia in BB25 represents the main difference with naive snail microbial community composition (B0/B25 vs BB25, student Τ test Ρ = 1.83 e -05) (Fig. 4B). We further observed decrease in Planctomycetes following sympatric primo-infection (B0/B25 vs BB1/BB4, student Τ test Ρ = 5.7 e-07) and a decrease in Cyanobacteria following allopatric primo-infection (B0/B25 vs BV1/BV4, student Τ test Ρ = 0.00254). Interestingly, the relative abundance of both phyla decreased following the challenge regardless of the challenge type (Planctomycetes B0/B25/BB25 vs BBB1/BBB4/BBV1/BBV4, student Τ test Ρ = 0.0017 and Cyanobacteria B0/B25/BB25 vs BBB1/BBB4/BBV1/BBV4, student Τ test Ρ = 0.000568) (Fig. 4B). Bacteroidetes decreased 25 days after sympatric primo-infection (B0/B25 vs BB25, student Τ test Ρ = 0.0467) but increased 4 days after heterologous challenge (BB25 vs BBV4, student Τ test Ρ = 0.00577). To conclude, depending on the phylum, we observed the largest modifications in microbiota composition at day 1 or 4 after primo-infections and challenges. Whereas the microbiota responded differently to sympatric and allopatric primo-infections, we found no significant differences between the homologous or heterologous challenge (see BBB1 vsBBV1, Fig. 4) (Table S5).

Finally, we examined the core microbiota dynamics during infection. Similarly, to the complete microbiota, the core-microbiota, consisting of 62 families, was affected by the type of infection (naive vs primo-infection (p=0.003) and naive ι/s challenge (p=0.003)), by the time of infection (early (1 day) vs late (4 days)) (p=0.006) and also by the strain of parasite used for primo-infection (SmBRE vs SmVEN) (p=0.016) (Table S6). Principal Coordinate Analysis (PCoA) based on Bray-Curtis dissimilarities between the core microbiotas (Fig. S1) yielded very similar results to those based on the entire dataset (Fig. 4A).

In addition, we observed that 69.4% (43) of the core microbiota families were significantly affected by infection. Among those, 6.5% (4 families) were affected regardless of the infection type (Table S7). Interestingly, these families belong to the most abundant ones (Table S8). Nineteen families (30.6%) were never affected by infection (Table S7). Those families belong to the less represented ones, with the exception of *Xanthomonadaceae,* which was the 7^th^ most represented family (see Table S8). Further 32.6% (14 families) of the core microbiota responded solely to the primo-infection and 4.7% (2 families) to the challenge (Table S7). Similarly, 20.9% (9 families) of the core microbiota was affected early after infection (1 day) and 7% later (3 families, day 4). Finally, 23.3% (10 families) were affected by the SmBRE infection and 9.3% (4 families) by the SmVEN infection (Table S7).

### Link between the microbiota dysbiosis and B. glabrata antimicrobial immune response

The expression level of transcripts encoding antimicrobial peptides and antimicrobial proteins (AMP) was investigated following sympatric or allopatric primo-infection and homologous or heterologous challenges using RNAseq data (Fig. 5 and Fig. S2). Two achacin genes, 5 lipopolysaccharide-binding protein / bactericidal permeability-increasing protein (LBP/PBI) genes, and 5 biomphamacin genes were selected based on the *Biomphalaria glabrata* genome annotation (Adema et al., 2017). In comparison with the naive snails, the LBP/BPI 3.1 and 3.2 were over-expressed at day 25 after infection and following the challenges, while all other genes of this family were under-expressed in all conditions (Fig. S2). The achacins were under-expressed following primo-infection in the sympatric combination only, and at day 25 after primo-infection and following both challenges (Fig. S2). No changes in expression were observed following allopatric primo-infection compared to the naive snails (Fig. S2). Finally, the AMP biomphamacins 1, 4, 5 and 6 were over-expressed throughout the infection stages, except for BV1 and BB25 (Fig.6). The biomphamacin 3 was mainly under-expressed except at BV4 and BBB, where no differential expressions have been detected compared with naive snails (Fig. 5).

**Fig. 5:**
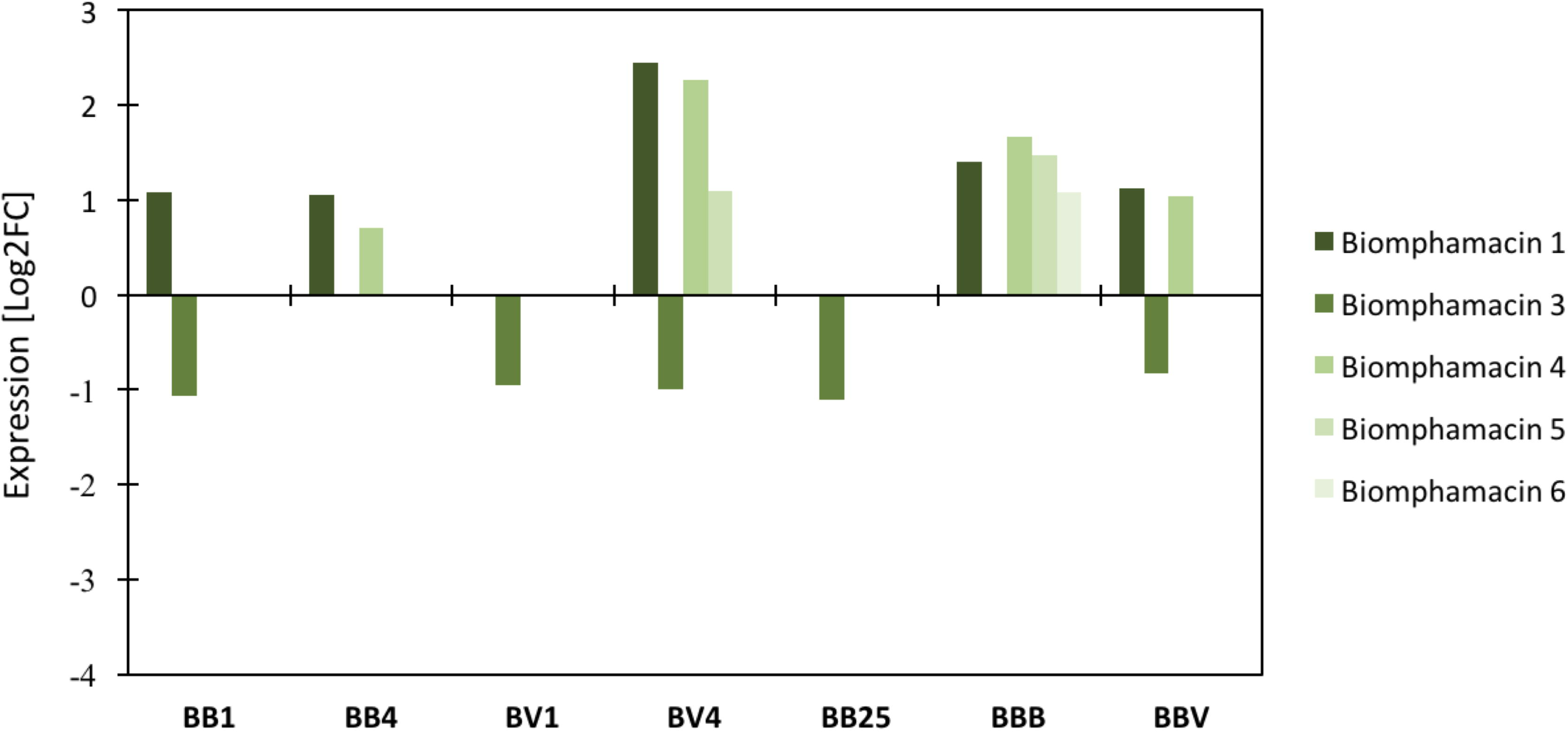
Differential gene expression of Biomphamacin antimicrobial peptides. Log2FC (fold change) of antimicrobial immune transcripts between naive and infected snails inferred from previous RNAseq analysis on the same experiment. A positive Log2 fold-change indicates over-expression in infected snails compared to the naive snails. Antimicrobial peptide families included 6 Biomphamacins (macin-like AMPs) consisting of 6 genes (shade of green).

Based on our observations of microbial community shifts, changes in expression of antimicrobial molecules were expected to occur following primo-infections and challenges and not for BB25, where resilience of microbial communities has been observed (Fig. 4 and Fig. S1). In this context, the LBP/BPI and achacin seem not to be linked with dysbiosis, as both remained highly down regulated even at BB25 when communities had already recovered (Fig. S2). All biomphamacins, except the biomphamacin 3, were over-expressed after infection, from day 1 in BB infection and day 4 in BV infection and for homologous or heterologous challenges (BBB and BBV). Finally, only a subset of biomphamacins (1, 4, 5, 6) changed in the course of infection but were not differentially expressed at BB25 compared with the naive snails (Fig. 5), suggesting their possible link with dysbiosis.

## Discussion

Fine-tuned interactions between microbiota, host immunity and pathogens have been observed in many vertebrate and invertebrate models (Rodrigues et al., 2010; Ramirez et al., 2012; Futo et al., 2015; Sansone et al., 2015; Chen et al., 2017). Following an infection by a pathogen, a host displays an innate or adaptive immune response against the intruder. The activation of such immune response may in turn affect the microbial community structure. Herein, we investigated the interactions between the host immune system, parasite and the bacterial microbiota in an invertebrate model - the gastropod snail *Biomphalaria glabrata and its* trematode parasite *Schistosoma mansoni.* Depending on the past evolutionary history between snails and schistosomes, different immune responses against 5. *mansoni* have been observed. We showed recently that in a sympatric interaction, the parasite that coevolved with its host induced a strong immunosuppression, whereas allopatric interaction resulted in a strong host cellular immune response (Portet et al., 2019). Moreover, a cellular immune response was observed following primo-infection, but a humoral immune response was observed following homologous or heterologous challenges (Pinaud et al., 2016). So using appropriate parasite-host combinations, we have the opportunity to modulate the host immune response type and to test changes in bacterial community composition and diversity (dysbiosis). First, studying the global bacterial microbiota community of *Biomphalaria* snails, we showed that the bacterial alpha diversity was not modified following primo-infection whatever the parasite strain or the time point of infection (Fig. 3, S3 Table). A decrease in alpha diversity is observed exclusively following challenge infections, as reflected by multiple indices (Fig. 3, S3 Table). Conversely, primo-infection and challenge strongly affected the bacterial OTU composition (Fig. 4). Moreover, differences in immunobiological interactions (immunosuppression, immune cellular response or immune humoral response) resulted in different microbiota dysbiosis reflected by specific changes in the snail microbial community (Fig. 4). Interestingly, homologous and heterologous challenges activated similar humoral immune response (Pinaud et al., 2016; Portet et al., 2019) that resulted in a similar change of microbiota alpha diversity and composition (Fig. 3, S3 Table, Fig. 4). Then, four days after infection, regardless of its type, the microbiota was still very different from the one in the naive snails, but the differences between the primo-infection and challenge disappeared, as apparent from the grouping of BB4, BV4, BBB4 and BBV4 in the PCoA (see Fig. 4). Furthermore, most of the OTUs affected by the primo-infection, returned to their initial state by the day 25, indicating that the snail microbiota is resilient against the infection (Fig. 4). However, some differences persisted: the Verrucomicrobia phylum remained highly represented in the infected snails at day 25 after primo-infection (Fig. 4B). Interestingly, some Verrucomicrobia species have been recently proposed as a hallmark of healthy gut due to its anti-inflammatory and immune-stimulant properties and its ability to improve gut barrier function (Fujio-Vejar et al., 2017). High abundance of Verrucomicrobia in the recovered snails could thus potentially reflect their role in community restoration (Fig. 4). Moreover, the expansion of a Verrucomicrobia species *Akkermansia muciniphilahas* been demonstrated in the gut of 5. *mansoni-*infected mice, suggesting a potentially functional role of Verrucomicrobia in *Schistosoma* infection processes in both final and intermediate hosts (Jenkins et al., 2018). We also paid a particular attention to the core-microbiota. Given the various ways to define core microbiota, we considered exclusively the persistent occurrence in the bacterial community in naive *Biomphalaria glabrata* snails (Astudillo-García et al., 2017). Similarly, to the entire microbiota, the core microbiota was affected by the type of infection (naive ι/s challenge), by the time of infection (day 1 vs day 4) as well as by the parasite strain (SmBRE vs SmVEN) (Table S6). Core microbiota seemed to be affected by immunosuppression or the activation of the immune response in a similar way than the whole microbiota (Fig. S1 and Fig. 4A). Understanding the shifts in core microbiota following infection is important as the long-term stability and persistent occurrence of beneficial microbes and their associated functions may contribute to host health and well-being to maintain host functionality and fitness toward changing ecological environment or environmental stress (Nyholm and McFall-Ngai, 2004; Lozupone et al., 2012; McFall-Ngai et al., 2013).

Given that infections affect the total and core microbiota structure, we explored the antimicrobial immune response following infection to find potential molecular mechanisms involved in the observed microbiota dysbiosis (Fig. 5 and Fig. S2). We analysed antimicrobial peptides (AMP) and antimicrobial proteins and our results suggest that only the AMP belonging to biomphamacin family may potentially affect the microbial communities in response to *Schistosoma* infection. Indeed, even though antimicrobial families can be involved in complex process of regulation or of control of microbiota communities, solely the biomphamacins AMP family members specifically displayed an expression pattern that can be linked with the observed dysbiosis of *Biomphalaria* bacterial communities (Fig. 5). Even if AMPs is often considered as the main immune pathway responsible for microbiota regulation (Onchuru and Kaltenpoth, 2019), an overall modification of the immune system may also be considered as a cause of dysbiosis. The potential role of immune recognition or ROS/NOS pathways as to be considered (Wang et al., 2014; Budachetri and Karim, 2015; Yang et al., 2015; Yang et al., 2016) and this will deserve further investigations in the present biological experimental model. It has been shown that the immune system is a key determinant of host-associated bacterial communities in many biological systems. Two mechanisms have been proposed to explain host-microbiota interactions through crosstalk with the host innate immune system. The first one proposes that the host immune system exerts constant pressure on the microbiota in order to maintain homeostasis (Hooper et al., 2012), the host immune system can thus control the composition of the resident microbiota (Zhang et al., 2015). According to this hypothesis, any changes in host immune response to infection would potentially affect the resident microbiota diversity and composition. The second mechanism proposes that the immune system would be tolerant to weak and continuous antigenic immune stimulations experienced during a lifespan (Pradeu and Eric, 2014), and thus that host immune system would not exert any pressure on the resident microbiota. In this case, any changes in host immune response to infection would potentially not affect the resident microbiota diversity and composition.

Interestingly, numerous studies demonstrate a direct control of microbial communities by the host immune system. As an example, species-specific antimicrobial peptides can shape species-specific bacterial associations in *Hydra* (Franzenburg et al., 2013). Other immune pathways have been also demonstrated to regulate or control the microbiota communities, like the intestinal homeo-box gene Caudal in *Drosophila* (Ryu et al., 2008), or neuropeptides with antibacterial activity which are secreted to shape the microbiome on the body surface of *Hydra* (Augustin et al., 2017), even host lectins were demonstrated to stabilize microbiota communities (Dinh et al., 2018).

Thus, if biotic stress (i.e. an infection) modifies the expression of antimicrobial peptides or other immune related pathways, an effect on the microbial communities can be expected. Our results indicate that bacterial communities could indeed be shaped by the immune system of B. *glabrata.* Based on the hypothesis proposed by Hooper and collaborators (Hooper and Macpherson, 2010; Hooper et al., 2012; Dinh et al., 2018), it seems that the immune system of *B. glabrata* snails maintains or controls the microbial communities permanently. Therefore, following an infection, the immune system is diverted from its function of managing the microbiota and consequently the bacterial communities evade its regulation, resulting in changes of their composition and diversity. When the immune response returns to a basal level, the microbiota is again under-control and the microbiota shows its resilience by returning to its “healthy” state (Hooper and Macpherson, 2010; Hooper et al., 2012), as observed in the present study (Fig. 4). In other words, the immune system is likely no longer able to maintain the microbiota homeostasis after infection (resulting in dysbiosis), which, may in turn affect host homeostasis or fitness (Contijoch et al., 2019; Warne et al., 2019) (Contijoch, Britton et al. 2019, Warne, Kirschman et al. 2019).

Changes in microbiota composition may result from shifts in the abundance of specific bacterial groups participating in anti-pathogen response, or just be a collateral effect of the immune response activation against metazoan parasite infection. This question will deserve further investigation, by testing 5. *mansoni* prevalence and intensity in experimental infections of *Biomphalaria glabrata* snails following antibiotic treatment or microbiota transplantation.

In order to fully understand Schistosomiasis transmission and to develop new ways to control the expansion of this widespread human parasitic disease in the field, it will be crucial to determine if the snail-associated bacterial communities affect the parasite transmission. For example, expanding knowledge on *Biomphalaria* snail microbiota is an essential step for developing paratransgenetic solutions to the spread of Schistosomiasis, involving the use of transgenic bacteria expressing foreign gene products (i.e., schistosomicidal compounds) that can reduce host competence and block pathogen development or transmission when introduced into the microbiota of vector snail field populations (Aksoy et al., 2008; Coutinho-Abreu et al., 2010; Gilbert et al., 2016).

### Experimental procedures

#### Ethical statements

Our laboratory holds permit # A66040 for experiments on animals, which was obtained from the French Ministry of Agriculture and Fisheries and the French Ministry of National Education, Research, and Technology. The housing, breeding and care of the utilized animals followed the ethical requirements of our country. The experimenter possesses an official certificate for animal experimentation from both of the above-listed French ministries (Decree # 87-848, October 19, 1987). The various protocols used in this study have been approved by the French veterinary agency of the DRAAF Languedoc-Roussillon (Direction Régionale de l’Alimentation, de l’Agriculture et de la Forét), Montpellier, France (authorization # 007083).

#### Biological material

In this study, we used a single strain of the host snail, the albino Brazilian strain of *Biomphalaria glabrata* (BgBRE) and two strains of the trematode parasite *Schistosoma mansoni:* a Brazilian strain (SmBRE, for sympatric infection) and a Venezuelan strain (SmVEN, for allopatric infection). BgBRE and SmBRE strains originate from the locality of Recife, Brazil; the SmVEN parasite strain was recovered from the locality of Guacara, Venezuela. All host and parasite strains were maintained in the laboratory on their respective sympatric snail hosts (SmBRE on BgBRE and SmVEN on BgVEN). The snails were reared at constant temperature of 26°C and fed only with washed salad every 3 days.

#### Experimental Infections (Fig. 1)

In order to decipher the inter-individual structure of snail microbiota as well as investigate the influence of different snail immune stimulations on its microbiota structure and dynamics, we applied a two-step experimental infection protocol (Fig. 1).

Briefly, BgBRE snails were primo-infected with one of the two parasite strains (SmBRE sympatric or SmVEN allopatric). The snails were then sampled 1 and 4 days following primo-infection by both parasite strains and 25 days after primo-infection for SmBRE infection condition. Then, the BgBRE snails primo-infected with SmBRE were challenged (25 days after the primo-infection) with SmBRE or SmVEN parasite strains for homologous or heterologous challenges respectively. The challenged snails were then sampled 1 and 4 days following challenge (Fig. 1). For all experimental infections, the snails were individually exposed for 12h to 10 miracidia in 5mL of pond water, thereafter snails were returned to the water tanks and separated according to the parasite strain and infection conditions.

##### Infections and sampling for microbiota analysis

We performed a primo-infection of 63 BgBRE snails with either SmBRE (sympatric, 49 snails) or SmVEN (allopatric, 14 snails) strains. Then, 25 days after the primo-infection, we challenged a subset (28 snails) of SmBRE-primo-infected snails with either SmBRE (14 snails) (homologous challenge) or SmVEN (14 snails) (heterologous challenge).

To examine the effect of primo-infection on the snail microbiota, we sampled 7 whole snails for each infection combination on day one (named BB1 and BV1; the first letter refers to the *Biomphalaria* strain BgBRE and the second letter refers to the origin of S. *mansoni* strains used for primo-infection, SmBRE or SmVEN), on day four (BB4 and BV4), and on day twenty-five (BB25, no SmVEN-infected snails were sampled on day 25) after the primo-infection. This BB25 sample is used as control for microbiota changes observed following challenge infections. Thus then, we sampled 7 whole-snails on day one after the challenge (BBB1 and BBV1; the third letter refers to the origin of the 5. *mansoni* strain used for the challenge, SmBRE or SmVEN) and on day four (BBB4 and BBV4) after the challenge. In addition, we used 6 naive snails collected at the beginning of the experiment (B0) and 6 naive snails collected at the time of challenge (i.e. day 25, named B25) as controls to assess the variability and stability of the BgBRE snail’s bacterial communities in our breeding and rearing laboratory conditions. Snails were not fed 24 hours before the DNA extraction for metabarcoding analysis.

##### Infection and sampling for host antimicrobial immune response

We performed a primo-infection of 180 BgBRE snails with either SmBRE (sympatric, 140 snails) or SmVEN (allopatric, 40 snails). Then, 25 days after infection, we challenged a subset (80 snails) of SmBRE-infected snails with either SmBRE (40 snails) (homologous challenge) or SmVEN (40 snails) (heterologous challenge) (Portet et al., 2019).

To study the primo-infection transcriptomic immune response of the snail, we took 20 snails pooled for each infection type on day one (BB1 and BV1), four (BB4 and BV4), and twenty-five (BB25). After the challenge with SmBRE or SmVEN, we sampled 20 snails in each condition at days 1 and 4. Then samples of day 1 and day 4 were mixed together into a single sample referring to BBB or BBV. In addition, we used 2 pools of 30 naive snails (B0.1 and B0.2) to establish the basal snail transcriptomic activity (Portet et al., 2019).

#### Extraction and sequencing

##### DNA extraction and 16S rDNA sequencing

Immediately after sampling, snail shells were cleaned with alcohol and removed, whole snails were then frozen in liquid nitrogen and grounded. The total DNA was extracted with DNeasy Blood and Tissue Kit (Qiagen) according to the manufacturer’s protocol. The DNA quantification was performed by Qubit 2.0 Fluorometer, using dsRNA BR Assay kit. Individual 16S rDNA amplicon libraries were generated using the 341F (CCTACGGGNGGCWGCAG) and 805R (GACTACΗVGGGΤATCTAATCC) primers targeting the variable V3-V4 loops (Klindworth et al., 2013). Paired-end sequencing with 250 bp read length was performed on the Illumina MiSeq sequencing system (Genome Québec, Montréal, Québec, Canada) using the v2 chemistry according to the manufacturer’s protocol.

##### RNA extraction and transcriptomic sequencing

Immediately after sampling, snail shells were cleaned with alcohol and removed, then snails were pooled according to infection type. Total RNA was extracted using TRIZOL (Sigma Life Science, USA) according to the manufacturer’s instructions. For BBB and BBV equimolar amounts of RNA extracted from molluscs challenged at both 1 and 4 days were mixed together into a single sample. mRNAs were sequenced in paired-end 72-bp read lengths, using the mRNA-Seq kit for transcriptome sequencing on Illumina Genome Analyzer II (MGX-Montpellier GenomiX, Montpellier, France).

#### Microbiota analysis

##### Data analysis of 16S sequences

The FROGS pipeline (Find Rapidly OTU with Galaxy Solution) implemented on a galaxy instance (http://sigenae-workbench.toulouse.inra.fr/galaxy/) was used for data processing (Escudié et al., 2017). Briefly, paired reads were merged using FLASH (Magoc and Salzberg, 2011). After denoising and primer/adapters removal (Didion et al., 2017), de novo clustering was done using SWARM, which uses a local clustering threshold, with aggregation distance d=3 after denoising. Chimeras were removed using VSEARCH (Rognes et al., 2016). We filtered out the singletons and performed taxonomic assignment using Blast+ against the Silva database (release 128).

All statistical analyses were done using R v3.3.1 (R: a language and environment for statistical computing, 2008; R Development Core Team, R Foundation for Statistical Computing, Vienna, Austria [http://www.R-project.org]). We used the phyloseq R package for community composition analysis (McMurdie and Holmes, 2013) to infer alpha diversity metrics as well as beta diversity (between sample distance). Beta diversity was examined by Principal Coordinate Analysis (PCoA) using the Bray-Curtis distance matrices. We performed a Mann Whitney U test (p<0.05) to compare alpha diversity and one-way PERMANOVA with a Benjamini & Hochberg post-hoc to analyze beta diversity between the experimental groups. For all analyses, the threshold significance level was set at 0.05.

##### Analysis of core-microbiota

We defined the core-microbiota as the set of bacterial families that were present in 100% of the naive individuals excluding unknown or multi-affiliations at lower taxonomic ranks (Table S1). We then used the abundances of OTUs belonging to these families to examine the composition of the core-microbiota.

To check if the core-microbiota is affected by infection, we compared the abundance of core families between infected snails and naive conditions with a one-way PERMANOVA with a Benjamini & Hochberg post-hoc correction. Moreover, a frequency test was performed to determine which specific families were affected during infection. The number of significantly differentially represented families at each sampling day (1, 4 and 25 days after primo infections and 1, 4 days after challenges) was calculated to assess the temporal variability during the course of infection.

#### Transcriptome analysis of antimicrobial immune response

##### Antimicrobial response

An antimicrobial transcriptome was built from transcripts known to be involved in *Biomphalaria* immune response against bacteria (antimicrobial peptides: biomphamacin or antimicrobial proteins: LBP/BPI and achacin; see Table S2 for details). The full-length sequences of these transcripts were recovered from GenBank and the *Biomphalaria* genome (Adema et al., 2017) and joined in a subset that represents the antimicrobial transcriptome of *B. glabrata.* This antimicrobial transcriptome was then concatenated with a de-novo assembled transcriptome of *Biomphalaria* available in our laboratory (see (Dheilly et al., 2015; Pinaud et al., 2016; Galinier et al., 2017) for details) and uploaded on the Galaxy server. Before concatenation, a blastn (70 % identity and 90% coverage) was conducted to identify redundant transcripts across the transcriptomes. Redundant transcripts were then discarded using CDhitEst to avoid mapping errors and bias in read counts when using Bowtie2 package.

##### Differential expression analysis

High-quality reads (Phred score >29) were aligned to the concatenated transcriptome using Bowtie2 (v.2.0.2), which was run locally on a Galaxy server. The DESeq2 (v2.12) R package was used to identify differential expression levels between uninfected (B0.1 and B0.2) and infected conditions (p value < 0.05).

## Acknowledgments

BG was supported by ANR JCJC INVIMORY (number ANR 13-JSV7-0009) from the French National Research Agency (ANR). The authors want to thanks the DHOF program of the UMR5244/IHPE (http://ihpe.univ-perp.fr/en/ihpe-transversal-holobiont/) for partial financialsupport. This study is set within the framework of the “Laboratoires d’Excellences (LABEX)” TULIP (ANR-10-LABX-41).

## Legends to Figures

Fig. SI: Core Microbiota analysis

Modifications of the bacterial core microbiota of *Biomphalaria glabrata* following *Schistosoma* primo-infection and challenge. After a one-way Permanova with a Benjamini & Hochberg post-hoc, a frequency test was realised to determine which specific families were affected during infection. A. Proportion of OTU Families affected by infection are represented by colour circles. The Venn diagram shows more precisely the specific impact of infections on the OTU families partially influenced. B. Heatmap corresponding to all core microbiota with clustering on Familiy modification patterns between conditions. The scale of red correspond to ratio between naive condition and each point of infection.

Fig. S2: Differential gene expression of LBP/BPI and Achacin antimicrobial protein families

Log2FC (fold change) of antimicrobial immune transcripts between naive and infected snails inferred from previous RNAseq analysis on the same experiment. A positive Log2 fold-change indicates over-expression in infected snails compared to the naive snails. Antimicrobial protein families: LBP/BPIs (*Lipopolysaccharide-binding protein / Bactericidal Permeability-Increasing protein)* consisting of 5 genes (in blue) and achacins consisting of 2 genes (in orange).

## Notes

#### Summary of Updates

We have greatly enhanced the analysis of the results and conclusions of the paper have changed.

## Reference

Adema, C.M., Hillier, L.W., Jones, C.S., Loker, E.S., Knight, M., Minx, P. et al. (2017) Whole genome analysis of a schistosomiasis-transmitting freshwater snail. Nature communications 8: 15451.

Aksoy, S., Weiss, B., and Attardo, G. (2008) Paratransgenesis applied for control of tsetse transmitted sleeping sickness. Adv Exp Med Biol 627: 35–48.

Allan, E.R.O., Tennessen, J.A., Sharpton, T.J., and Blouin, M.S. (2018) Allelic Variation in a Single Genomic Region Alters the Microbiome of the Snail Biomphalaria glabrata. J Hered 109: 604–609

Astudillo-García, C., Bell James, J., Webster Nicole, S., Glasl, B., Jompa, J., Montoya Jose, M., and Taylor Michael, W. (2017) Evaluating the core microbiota in complex communities: A systematic investigation. Environmental Microbiology 19:1450–1462.

Augustin, R., Fraune, S., and Bosch, T.C.G. (2010) How Hydra senses and destroys microbes. Seminars in Immunology 22: 54–58.

Augustin, R., Schröder, K., Murillo Rincón, A.P., Fraune, S., Anton-Erxleben, F., Herbst, E.-M. et al. (2017) A secreted antibacterial neuropeptide shapes the microbiome of Hydra. Nature communications 8: 698.

Bosch, T.C. (2013) Cnidariari-microbe interactions and the origin of innate immunity in metazoans. Annu Rev Microbiol 67: 499–518.

Bouchut, Α., Roger, E., Gourbal, B., Grunau, C., Coustau, C., and Mitta, G. (2008) The compatibility polymorphism in invertebrate host/trematode interactions: research of molecular determinants. Parasite 15: 304–309.

Budachetri, K., and Karim, S. (2015) An insight into the functional role of thioredoxin reductase, a selenoprotein, in maintaining normal native microbiota in the Gulf Coast tick (Amblyomma maculatum). Insect Mol Biol 24: 570–581.

Chen, F., Krasity, B.C., Peyer, S.M., Koehler, S., Ruby, E.G., Zhang, X., and McFall-Ngai, M.J. (2017) Bactericidal Permeability-Increasing Proteins Shape Host-Microbe Interactions. mS/’o 8: e00040–00017.

Cirimotich, C.M., Dong, Y., Clayton, A.M., Sandiford, S.L., Souza-Neto, J.A., Mulenga, Μ., and Dimopoulos, G. (2011) Natural microbe - mediated refractoriness to Plasmodium infection in Anopheles gambiae. Science 332: 855–858.

Contijoch, E.J., Britton, G.J., Yang, C., Mogno, I., Li, Z., Ng, R. et al. (2019) Gut microbiota density influences host physiology and is shaped by host and microbial factors. Elife 8: pii: e40553.

Coustau, C., Gourbal, B., Duval, D., Yoshino, T.P., Adema, C.M., and Mitta, G. (2015) Advances in gastropod immunity from the study of the interaction between the snail Biomphalaria glabrata and its parasites: A review of research progress over the last decade. Fish & Shellfish Immunology 46: 5–16.

Coutinho-Abreu, I.V., Zhu, K.Y., and Ramalho-Ortigao, M. (2010) Transgenesis and paratransgenesis to control insect-borne diseases: current status and future challenges. Parasitol Int 59:1–8.

Dheilly, Ν.Μ., Duval, D., Mouahid, G., Emaris, R., Allierine, J.-F., Galinier, R. et al. (2015) A family of variable immunoglobulin and lectin domain containing molecules in the snail Biomphalaria glabrata. Developmental & Comparative Immunology 48: 234–243.

Didion, J.P., Martin, M., and Collins, F.S. (2017) Atropos: specific, sensitive, and speedy trimming of sequencing reads. PeerJ 5: e3720.

Dinh, C., Farinholt, T., Hirose, S., Zhuchenko, O., and Kuspa, A. (2018) Lectins modulate the microbiota of social amoebae. Science 361: 402–406.

Doenhoff, M.J., Hagan, P., Cioli, D., Southgate, V., Pica-Mattoccia, L., Botros, S. et al. (2009) Praziquantel: its use in control of schistosomiasis in sub-Saharan Africa and current research needs. Parasitology 136:1825–1835.

Ducklow, H.W., Boyle, P.J., Maugel, P.W., Strong, C., and Mitchell, R. (1979) Bacterial flora of the schistosome vector snail Biomphalaria glabrata. Applied and Environmental Microbiology 38: 667–672.

Escudié, F., Auer, L., Bernard, M., Mariadassou, M., Cauquil, L., Vidal, Κ., et al. (2017) FROGS: Find, Rapidly, OTUs with Galaxy Solution. Bioinformatics: btx791.

Fallon, P.G., and Doenhoff, M.J. (1994) Drug-Resistant Schistosomiasis: Resistance to Praziquantel and Oxamniquine Induced in Schistosoma Mansoni in Mice is Drug Specific. The American Journal of Tropical Medicine and Hygiene 51: 83–88.

Franzenburg, S., Fraune, S., Künzel, S., Baines, J.F., Domazet-Lošo, T., and Bosch, T.C.G. (2012) MyD88-deficient Hydra reveal an ancient function of TLR signaling in sensing bacterial colonizers. Proceedings of the National Academy of Sciences of the United States of America 109: 19374–19379.

Franzenburg, S., Walter, J., Kürizel, S., Wang, J., Bairies, J.F., Bosch, T.C.G., and Fraune, S. (2013) Distinct antimicrobial peptide expression determines host species-specific bacterial associations. Proceedings of the National Academy of Sciences of the United States of America 110: E3730–E3738.

Fraune, S., Augustin, R., Anton-Erxleben, F., Wittlieb, J., Gelhaus, C., Klimovich, V.B. et al. (2010) In an early branching metazoan, bacterial colonization of the embryo is controlled by maternal antimicrobial peptides. Proceedings of the National Academy of Sciences of the United States of America 107: 18067–18072.

Fujio-Vejar, S., Vasquez, Y., Morales, P., Magne, F., Vera-Wolf, P., Ugalde, J.A. et al. (2017) The Gut Microbiota of Healthy Chilean Subjects Reveals a High Abundance of the Phylum Verrucomicrobia. Front Microbiol 8: 1221.

Futo, Μ., Armitage, S.A., and Kurtz, J. (2015) Microbiota Plays a Role in Oral Immune Priming in Tribolium castaneum. Front Microbiol 6: 1383.

Galinier, R., Roger, E., Moné, Y., Duval, D., Portet, Α., Pinaud, S., et al. (2017) A multistrain approach to studying the mechanisms underlying compatibility in the interaction between Biomphalaria glabrata and Schistosoma mansoni. PLOS Neglected Tropical Diseases 11: e0005398.

Gendrin, M. (2017) A Swiss Army Knife to Cut Malaria Transmission. Cell Host & Microbe 22: 577–579.

Gilbert, J.A., Medlock, J., Townsend, J.P., Aksoy, S., Ndeffo Mbah, M., and Galvani, A.P. (2016) Determinants of Human African Trypanosomiasis Elimination via Paratransgenesis. PLoS Negl Trop Dis 10: e0004465.

Hooper, L.V., and Macpherson, A.J. (2010) Immune adaptations that maintain homeostasis with the intestinal microbiota. Nat Rev Immunol 10:159–169.

Hooper, L.V., Littman, D.R., and Macpherson, A.J. (2012) Interactions between the microbiota and the immune system. Science (New York, N.Y.) 336:1268–1273.

Jenkins, T.P., Peachey, L.E., Ajami, N.J., MacDonald, A.S., Hsieh, M.H., Brindley, P.J. et al. (2018) Schistosoma mansoni infection is associated with quantitative and qualitative modifications of the mammalian intestinal microbiota. Sci Rep 8:12072.

Joshi, D., Pan, X., McFadden, M.J., Bevins, D., Liang, X., Lu, P. et al. (2017) The Maternally Inheritable Wolbachia wAIbB Induces Refractoriness to Plasmodium berghei in Anopheles stephensi. Frontiers in Microbiology 8: 366.

Klindworth, Α., Pruesse, E., Schweer, T., Peplies, J., Quast, C., Horn, M., and Glockner, F.O. (2013) Evaluation of general 16S ribosomal RNA gene PCR primers for classical and next-generation sequencing-based diversity studies. Nucleic Acids Res 41: el.

Lozupone, C.A., Stombaugh, J.I., Gordon, J.I., Jansson, J.K., and Knight, R. (2012) Diversity, stability and resilience of the human gut microbiota. Nature 489: 220–230.

Magoc, T., and Salzberg, S.L. (2011) FLASH: fast length adjustment of short reads to improve genome assemblies. Bioinformatics 27: 2957–2963.

McFall-Ngai, M., Hadfield, M.G., Bosch, T.C., Carey, H.V., Domazet-Loso, T., Douglas, A.E. et al. (2013) Animals in a bacterial world, a new imperative for the life sciences. Proc Natl Acad Sci U S A 110: 3229–3236.

McMurdie, P.J., and Holmes, S. (2013) phyloseq: an R package for reproducible interactive analysis and graphics of microbiome census data. PLoS One 8: e61217.

Mitta, G., Adema, C.M., Gourbal, Β., Loker, E.S., and Theron, A. (2012) Compatibility polymorphism in snail/schistosome interactions: From field to theory to molecular mechanisms. Dev Comp Immunol 37:1–8.

Mitta, G., Gourbal, B., Grunau, C., Knight, M., Bridger, J.M., and Théron, A. (2017) Chapter Three - The Compatibility Between Biomphalaria glabrata Snails and Schistosoma mansoni: An Increasingly Complex Puzzle. Advances in Parasitology 97:111–145.

Mitta, G., Galinier, R., Tisseyre, P., Allienne, J.F., Girerd-Chambaz, Y., Guillou, F. et al. (2005) Gene discovery and expression analysis of immune-relevant genes from Biomphalaria glabrata *hemocytes*. Developmental and comparative immunology 29: 393-407.

Mone, Y., Gourbal, B., Duval, D., Du Pasquier, L., Kieffer-Jaquinod, S., and Mitta, G. (2010) A large repertoire of parasite epitopes matched by a large repertoire of host immune receptors in an invertebrate host/parasite model. PLoS Negl Trop Dis 4: pii: e813.

Moné, Y., Ribou, A.-C., Cosseau, C., Duval, D., Théron, Α., Mitta, G., and Gourbal, B. (2011) An example of molecular co-evolution: Reactive oxygen species (ROS) and ROS scavenger levels in Schistosoma mansoni/Biomphalaria glabrata interactions. International Journal for Parasitology 41: 721–730.

Narasimhan, S., Schuijt, T.J., Abraham, N.M., Rajeevan, N., Coumou, J., Graham, M. et al. (2017) Modulation of the tick gut milieu by a secreted tick protein favors Borrelia burgdorferi colonization. Nature communications 8:184.

Nyholm, S.V., and McFall-Ngai, M.J. (2004) The winnowing: establishing the squid-vibrio symbiosis. Nat Rev Microbiol 2: 632–642.

Onchuru, T.O., and Kaltenpoth, M. (2019) Established cotton stainer gut bacterial mutualists evade regulation by host antimicrobial peptides. Appl Environ Microbiol pii: AEM.00738-19..

Pinaud, S., Pórtela, J., Duval, D., Nowacki, F.C., Olive, M.-A., Allienne, J.-F. et al. (2016) A Shift from Cellular to Humoral Responses Contributes to Innate Immune Memory in the Vector Snail Biomphalaria glabrata. PLOS Pathogens 12: el005361.

Portet, Α., Pinaud, S., Chaparro, C., Galinier, R., Dheilly, N.M., Pórtela, J. et al. (2019) Sympatric versus allopatric evolutionary contexts shape differential immune response in Biomphalaria / Schistosoma interaction. PLoS Pathog 15: el007647.

Pradeu, T., and Eric, V. (2014) The discontinuity theory of immunity. Sci Immunol 1: 1–9.

Ramirez, J.L., Souza-Neto, J., Cosme, R.T., Rovira, J., Ortiz, Α., Pascale, J.M., and Dimopoulos, G. (2012) Reciprocal tripartite interactions between the Aedes aegypti midgut microbiota, innate immune system and dengue virus influences vector competence. PLOS Neglected Tropical Diseases 6:1–11.

Rodrigues, J., Brayner, F.A., Alves, L.C., Dixit, R., and Barillas-mury, C. (2010) Hemocyte Differentiation Mediates Innate Immune Memory in Anopheles gambiae Mosquitoes. Science 329: 1353–1355.

Rognes, T., Flouri, T., Nichols, B., Quince, C., and Mahe, F. (2016) VSEARCH: a versatile open source tool for metagenomics. PeerJ 4: e2584.

Ryu, J.-H., Kim, S.-H., Lee, H.-Y., Bai, J.Y., Nam, Y.-D., Bae, J.-W. et al. (2008) Innate Immune Homeostasis by the Homeobox Gene Caudal and Commensal-Gut Mutualism in Drosophila. Science 319: 777.

Sansone, C.L., Cohen, J., Yasunaga, Α., Xu, J., Osborn, G., Subramanian, H., et al. (2015) Microbiota-dependent priming of antiviral intestinal immunity in Drosophila. Cell Host and Microbe 18: 571–581.

Silva, T.M., Meló, E.S., Lopes, A.C.S., Veras, D.L., Duarte, C.R., Alves, L.C., and Brayrier, F.A. (2013) Characterization of the bacterial microbiota of Biomphalaria glabrata (Say, 1818) (Mollusca: Gastropoda) from Brazil. Letters in Applied Microbiology 57:19–25.

Tennessen, J.A., Théron, Α., Marine, Μ., Yeh, J.-Y., Rognon, Α., and Blouin, M.S. (2015) Hyperdiverse Gene Cluster in Snail Host Conveys Resistance to Human Schistosome Parasites. PLoS Genetics 11: el005067.

Wang, X.W., Xu, J.D., Zhao, X.F., Vasta, G.R., and Wang, J.X. (2014) A shrimp C-type lectin inhibits proliferation of the hemolymph microbiota by maintaining the expression of antimicrobial peptides. J Biol Chem 289:11779–11790.

Warne, R.W., Kirschman, L., and Zeglin, L. (2019) Manipulation of gut microbiota during critical developmental windows affects host physiological performance and disease susceptibility across ontogeny. J Anim Ecol Epub ahead of print.

WHO (2002) TDR Strategic Direction for Research: Schistosomiasis. World Health Organization Information.

Xi, Z., Ramirez, J.L., and Dimopoulos, G. (2008) The Aedes aegypti toll pathway controls dengue virus infection. PLOS Pathogens 4(7): el000098.

Yang, H.T., Yang, M.C., Sun, J.J., Shi, X.Z., Zhao, X.F., and Wang, J.X. (2016) Dual oxidases participate in the regulation of intestinal microbiotic homeostasis in the kuruma shrimp Marsupenaeus japonicus. Dev Comp Immunol 59: 153–163.

Yang, H.T., Yang, M.C., Sun, J.J., Guo, F., Lan, J.F., Wang, X.W. et al. (2015) Catalase eliminates reactive oxygen species and influences the intestinal microbiota of shrimp. Fish Shellfish Immunol 47: 63–73.

Zhang, Η., Sparks, J.Β., Karyala, S.V., Settlage, R., and Luo, X.M. (2015) Host adaptive immunity alters gut microbiota. ISME J 9: 770–781.

