## Supplementary figures and images for "Infection of the *Biomphalaria glabrata vector snail* by *Schistosoma mansoni* parasites drives snail microbiota dysbiosis"

### Supplementary Figure 1

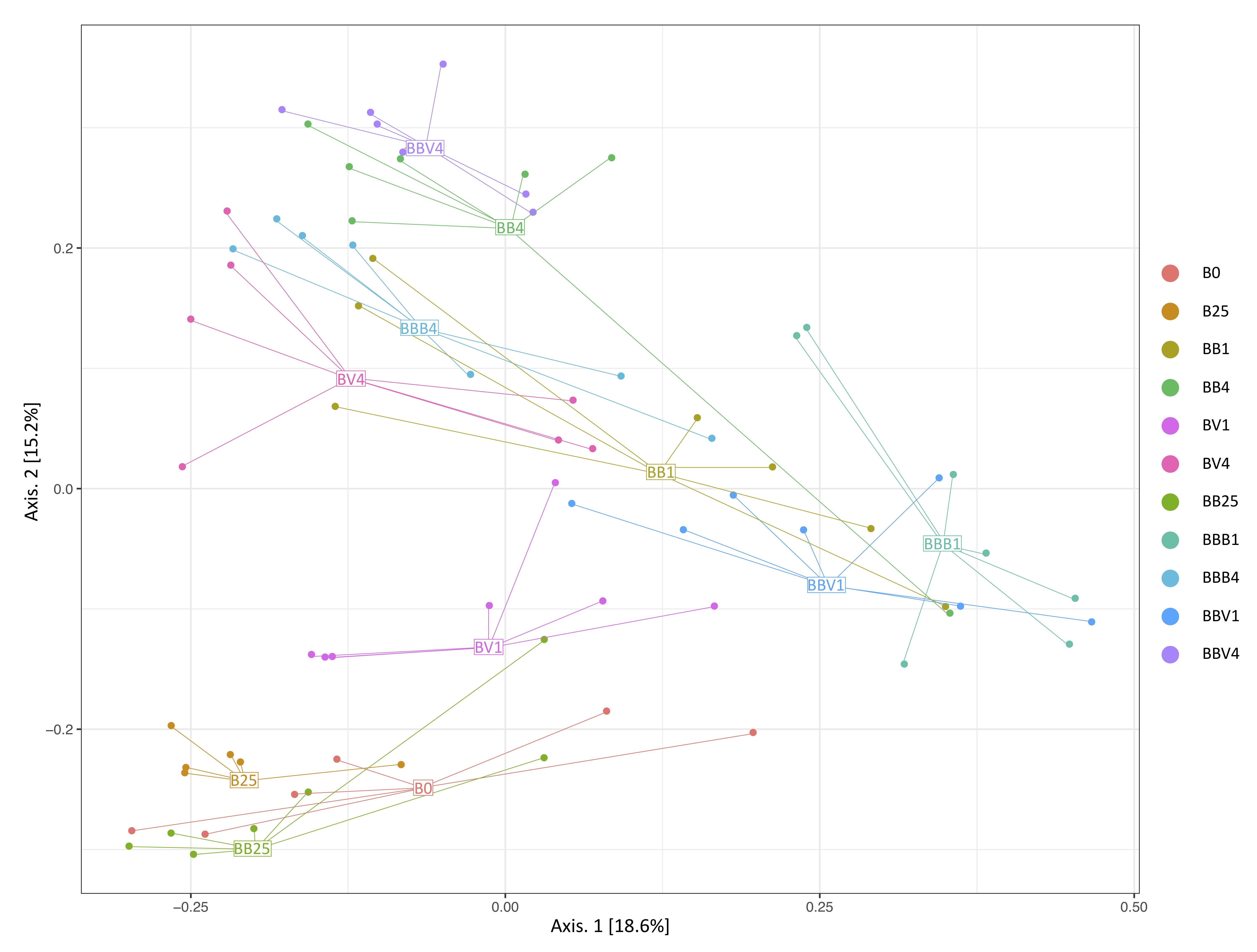

### Supplementary figure 2

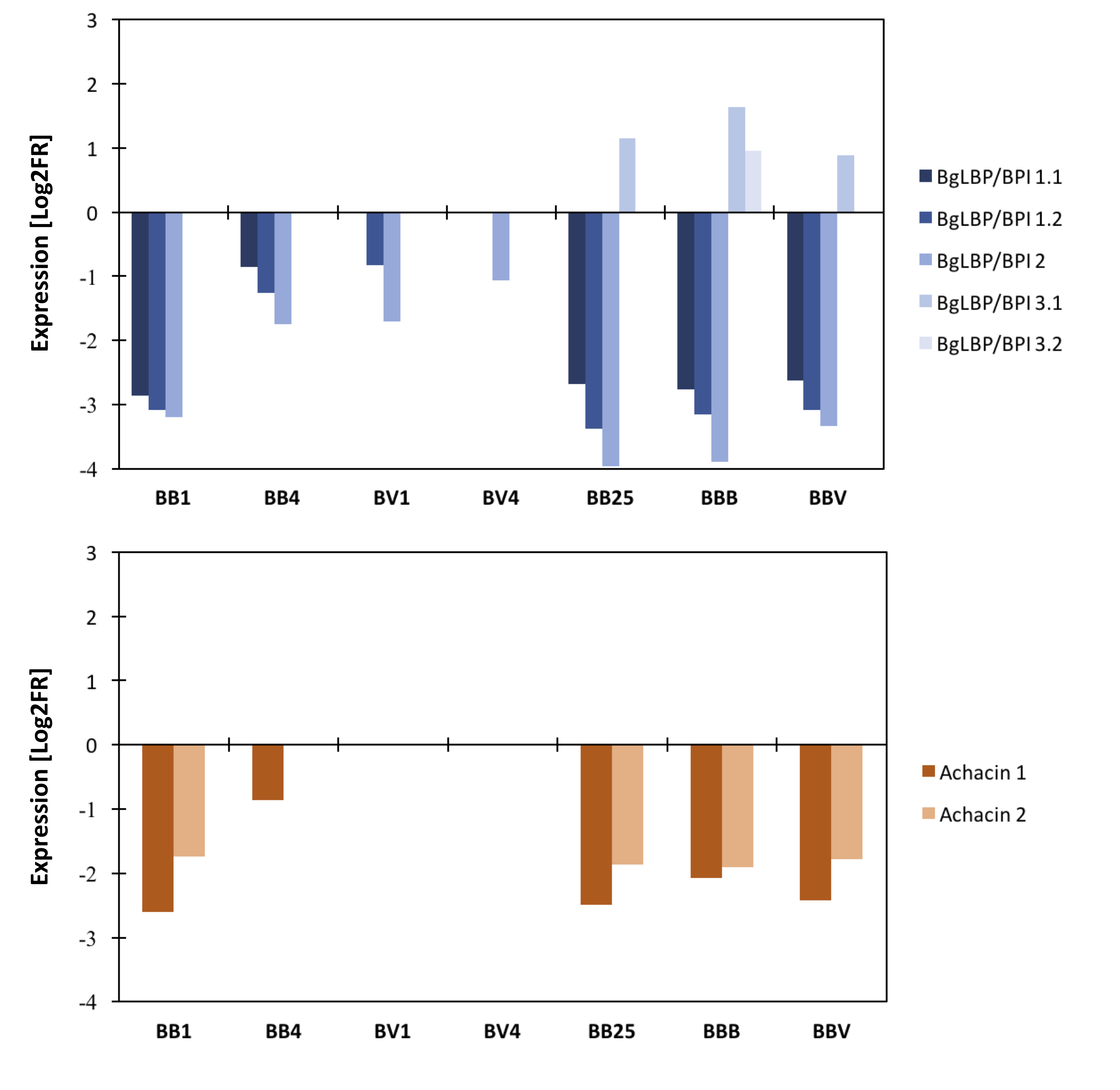
